# Spatial transcriptomics map of the embryonic mouse brain: a tool to explore neurogenesis

**DOI:** 10.1101/2023.08.10.552773

**Authors:** Barbara Di Marco, Javier Vázquez-Marín, Hannah Monyer, Lázaro Centanin, Julieta Alfonso

## Abstract

The developing brain has a well-organized anatomical structure comprising different types of neural and non-neural cells. Stem cells, progenitors, and newborn neurons tightly interact with their neighbouring cells and tissue microenvironment, and this intricate interplay ultimately shapes the output of neurogenesis. Given the relevance of spatial cues during brain development, we acknowledge the necessity for a spatial transcriptomics map accessible to the neurodevelopmental community. To fulfil this need, we generated spatially-resolved RNAseq data from E13.5 mouse brain sections immunostained for mitotic active neural and vascular cells. Unsupervised clustering defined specific cell type populations of diverse lineages and differentiation states. Differential expression analysis revealed unique transcriptional signatures across specific brain areas, uncovering novel features inherent to particular anatomical domains. Finally, we integrated existing single-cell RNAseq datasets into our Spatial Transcriptomics map, adding tissue context to single-cell RNAseq data. In summary, we provide a valuable tool that enables the exploration and discovery of unforeseen molecular players involved in neurogenesis, particularly in the crosstalk between different cell types.

**Summary Statement:** Di Marco, Vázquez-Marín et al. provide an open-access spatial transcriptomics atlas of the embryonic mouse brain. This spatial map enables the exploration and discovery of gene functions within tissue context.

## INTRODUCTION

The elaborated architecture of the mammalian brain relies on the perfect assembly of neuronal networks. The formation of these circuits is defined during embryonic development, a process strongly influenced by cues from the surrounding environment. Already from early neurogenic stages onwards, the developing brain exhibits a distinct compartmentalization in its anatomical structure. The dorsal and ventral germinal zones of the telencephalon consist of two proliferative areas populated by different cell types. Lining the wall of the cerebral ventricles sits the ventricular zone (VZ). The VZ hosts embryonic neural stem cells (NSCs), also known as radial glia cells, which have self-renewal capacity and the potential to give rise to neurons and glial cells. Adjacent to the VZ lies the subventricular zone (SVZ), an area more prominent in the ventral telencephalon. The SVZ is populated by intermediate progenitors that undergo several rounds of division and differentiation phases to produce young neurons (Taverna et al., 2014; Turrero García and Harwell, 2017). While NSCs and progenitors remain in the VZ/SVZ, newborn cortical neurons migrate away from the proliferative zones towards the pial surface, ultimately reaching either the preplate at early stages or the cortical plate at later embryonic ages (Hatanaka et al., 2016). To arrive at their final location, postmitotic cortical neurons follow different migratory routes guided by external cues (Marín et al., 2010). Subsequently, further maturation processes, such as dendrite and axon formation, synaptogenesis, and neuronal wiring, take place before the completion of brain development (Munno and Syed, 2003).

Advances in single-cell transcriptomics allowed the characterization of several cell types in the developing brain and helped to understand their functions (Chen et al., 2017; Moreau et al., 2021). However, single-cell transcriptomics lacks the spatial organization of neural and non-neural cells in the brain tissue, and the interconnection among different cell types is not preserved. This is a major drawback, given the functional significance of the tissue microenvironment during neurogenesis (Di Marco et al., 2020; Lange et al., 2016; Tan et al., 2016). Spatial transcriptomics (ST) technology has helped to surpass this limitation by assigning molecular profiles to specific locations within the tissue. Here, we launch an open-access dataset to browse gene expression data from mouse embryonic brain sections at the peak of neurogenesis. Importantly, our approach allows quantitative detection of mRNA in brain sections immunostained with markers for mitotic cells, blood vessels and radial glia cells. We show that ST unsupervised clustering defined specific embryonic brain areas according to their molecular hallmarks and revealed previously-unknown marker genes for each compartment. In addition, we present examples of manual clustering, a method that allows gene expression comparisons within custom-selected brain areas. Finally, we highlight ST and single-cell RNAseq (scRNAseq) data integration as a helpful approach to add tissue context to single-cell data.

## RESULTS AND DISCUSSION

### Spatial Transcriptomic in the embryonic mouse brain

We generated RNAseq data from embryonic mouse brains using ST. As a first step, we cryosectioned E13.5 heads from four wild-type embryos and collected the sections on Visium spatial gene expression slides (10x Genomics) (Fig. 1A). We used coronal sections containing the brain at the level of the lateral ganglionic eminences (LGE), a major neurogenic niche for GABAergic neurons and the source of adult neural stem cells (Fuentealba et al., 2015; Stenman et al., 2002). After fixation, sections were processed for immunofluorescent staining and imaging. We used Nestin to identify radial glia cells, CD31 to visualize vessels, and Phospho-Histone 3 (PH3) to label mitotic cells (Fig. 1B). To capture gene expression, the tissue was subsequently processed for library construction and sequencing (Fig. 1C). mRNA was released from the tissue and bound to spatially barcoded capture probes present in each spot (55 μm diameter area). Spatially defined cDNA libraries were synthesized and sequenced. Finally, gene expression values were mapped to the corresponding spots of the capture area by 10x Genomics Space Ranger software. We obtained data from 472 spots on average per brain, 61.676-105.113 mean reads per spot, and 3.262-3.815 median genes per spot.

**Figure 1.**
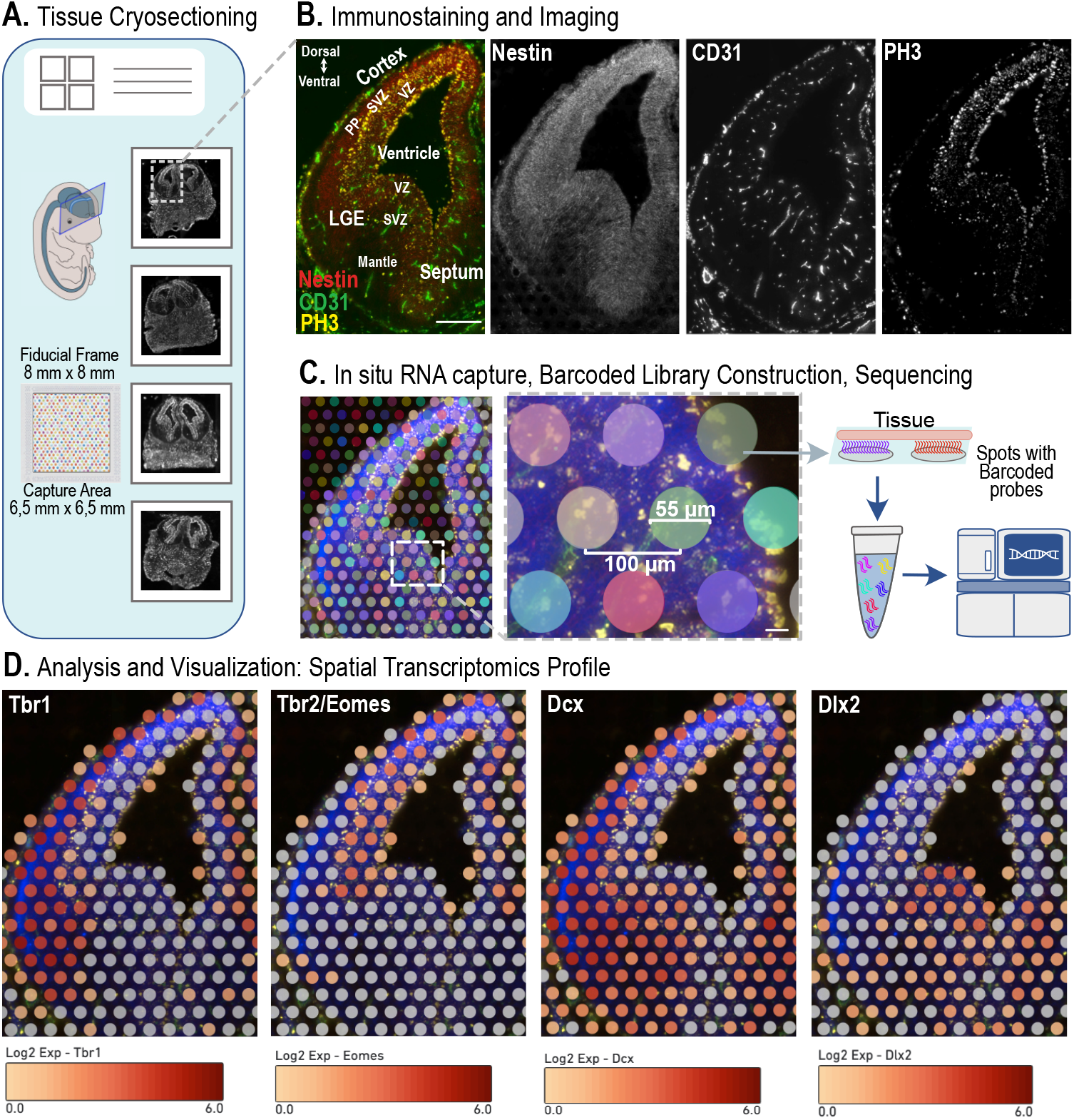
Spatial Transcriptomics workflow. **A**. E13.5 mouse embryonic brains were fresh frozen and cryosectioned. Brain sections were collected onto a spatial gene expression slide. **B**. Brain sections were fixed and processed for immunofluorescence staining and imaging. Nestin (radial glia cells, red), CD31 (endothelial cells and vessels, green), and PH3 (mitotic neural progenitors, yellow) were used as reference markers. LGE: lateral ganglionic eminence, VZ: ventricular zone, SVZ: subventricular zone, PP: preplate. Scale bar 200 μm. **C**. Tissue sections were permeabilized and processed for library construction. Each spot contains barcoded capture probes. Spatially defined cDNA libraries were sequenced. Scale bar 20 μm. **D**. Brain maps showing spatially resolved expression profiles for the indicated genes. Tissue images and sequencing data reconstruction were performed by Space Ranger software. Data visualization as a spatial gene expression map was obtained with Loupe Browser. Log2 scale illustrates colour-coded gene expression levels.

We visualized and investigated gene expression with 10x Genomics Loupe Browser, simultaneously detecting immunostainings and gene expression in the same brain section. As examples, we report spatially resolved expression profiles for *Tbr1, Tbr2/Eomes, Dcx*, and *Dlx2* (Fig. 1D). Importantly, we offer the spatial molecular atlas of the embryonic mouse brain readily available to the scientific community as an open-access resource at STOmicsDB (https://db.cngb.org/stomics/, Xu et al., 2022).

### Unsupervised clustering reveals spatially defined cell populations and specific marker genes

Unsupervised clustering of our RNAseq dataset from the four brain sections resulted in six clusters with a well-defined anatomical identity, reflecting the cellular organization of the developing brain (Fig. 2A-C). Indeed, unsupervised clustering faithfully reflected the main anatomical regions of the dorsal and ventral embryonic brain, the birthplace of glutamatergic and GABAergic neurons, respectively (Marín and Müller, 2014). Specifically, we found that Cluster 1 included the LGE SVZ and mantle zone, Cluster 2 the septal area, Cluster 3 the cortical SVZ and preplate, Cluster 4 the pial surface, Cluster 5 the cortical VZ, and Cluster 6 the LGE VZ (Fig. 2B-C). Besides gene expression visualization, we could investigate global differences in gene expression within the six brain clusters (Table S1). Among the top ten differentially-expressed genes (DEG), we found known markers for the different areas of the telencephalon. For instance, we identified *Neurog2* for the cortical VZ, *Ascl1* for the LGE VZ, *Neurod6* for the cortical SVZ/preplate, *Gad2* and *Dlx6* for the LGE SVZ, and *Zic* 4 and 5 for the septum. In addition, to validate the cluster identity with well-known molecular markers, we could assign a cluster-enriched expression to several other genes that were not previously recognized as markers of these particular brain areas. For example, *Sox3* for the cortical VZ, *Ddah1* for the LGE VZ, *Tiam2* for the cortical SVZ/preplate, *Tac1* for the LGE SVZ, and *Onecut2* and *Ecel1* for the septum (Fig. 2D). Of note, we could also detect gene expression signatures of non-neural populations in cluster 4. The Pia cluster covered a tissue area populated by fibroblasts and CD31+ pial vessels surrounding the hemispheres. Thus, the top ten DEGs encoded fibroblast-specific proteins (i.e., *Dcn* and *Col1a2*), but there were also genes enriched in endothelial cells (*Cldn5, Kdr*), blood cells (*Hba-x*) and microglia (*S100a11, Crybb1*) within the complete list (Fig. 2D and Table S1). Finally, we mapped gene expression levels for several cluster hallmarks to further confirm the accuracy of the clusters (Fig. 2E). In summary, our DEG analysis provides an exhaustive list of markers per brain area and helps elucidate the expression of several genes in specific cell populations according to their location.

**Figure 2.**
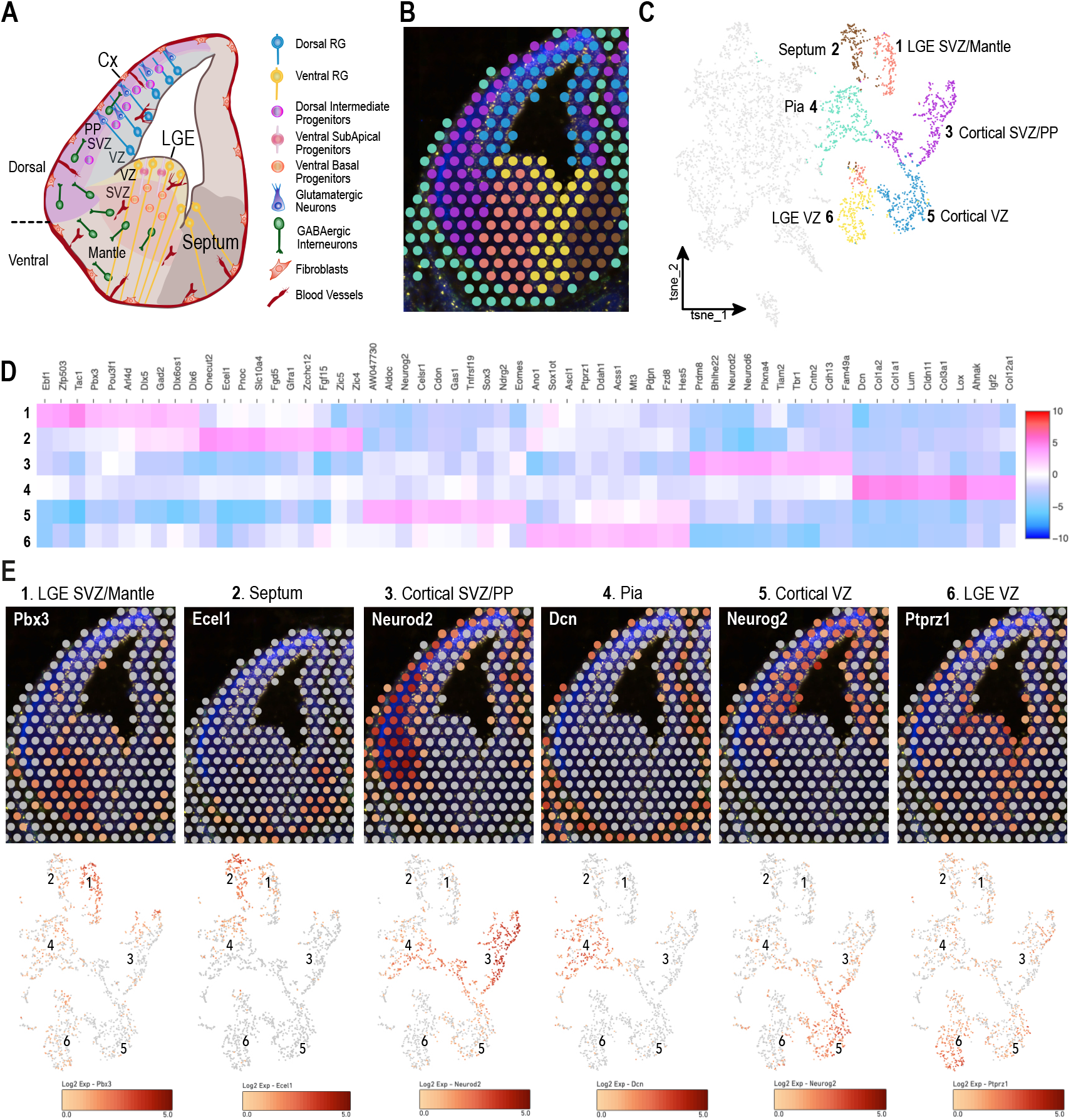
Unsupervised clustering of spatial transcriptomics data reveals spatially-segregated cell populations and specific marker genes. **A**. Schematic drawing depicting the main anatomical regions and cell populations of the dorsal and ventral embryonic brain at E13.5. **B**. Unsupervised clustering based on molecular signatures. Colour-coded clusters map into specific brain areas. **C**. Unsupervised clustering illustrated as a t-SNE plot. Cluster identity was assigned based on their anatomical location. **D**. Heatmap showing the top 10 differentially-expressed genes per cluster. **E**. Top: Gene expression visualization of representative cluster gene markers on a spatial map. Bottom: Feature plots showing the expression of the respective genes on t-SNE space.

### Spatial Transcriptomics enables manual cluster selection and comparison

To further exploit the potential of our newly generated database, we employed yet another analytic approach using manual cluster selection. We were able to select specific tissue areas based on the brain anatomy, as the original images for PH3, Nestin and CD31 immunostainings were aligned to the gene expression maps. In the first example, we compared the dorsal versus ventral telencephalon by manually selecting the corresponding spots. The dorsal cluster corresponded to the complete neocortical area, and the ventral cluster to the whole LGE area (Fig. 3A). Among the top upregulated genes, we found several involved in glutamatergic or GABAergic lineage specification, as expected from previous work (Marín and Müller, 2014). Additionally, using our DEG analysis we were able to uncover other genes newly identified as highly enriched in these specific areas, for instance, *Rprm* in the neocortex and *Carhsp1* in the LGE (Table S2).

**Figure 3.**
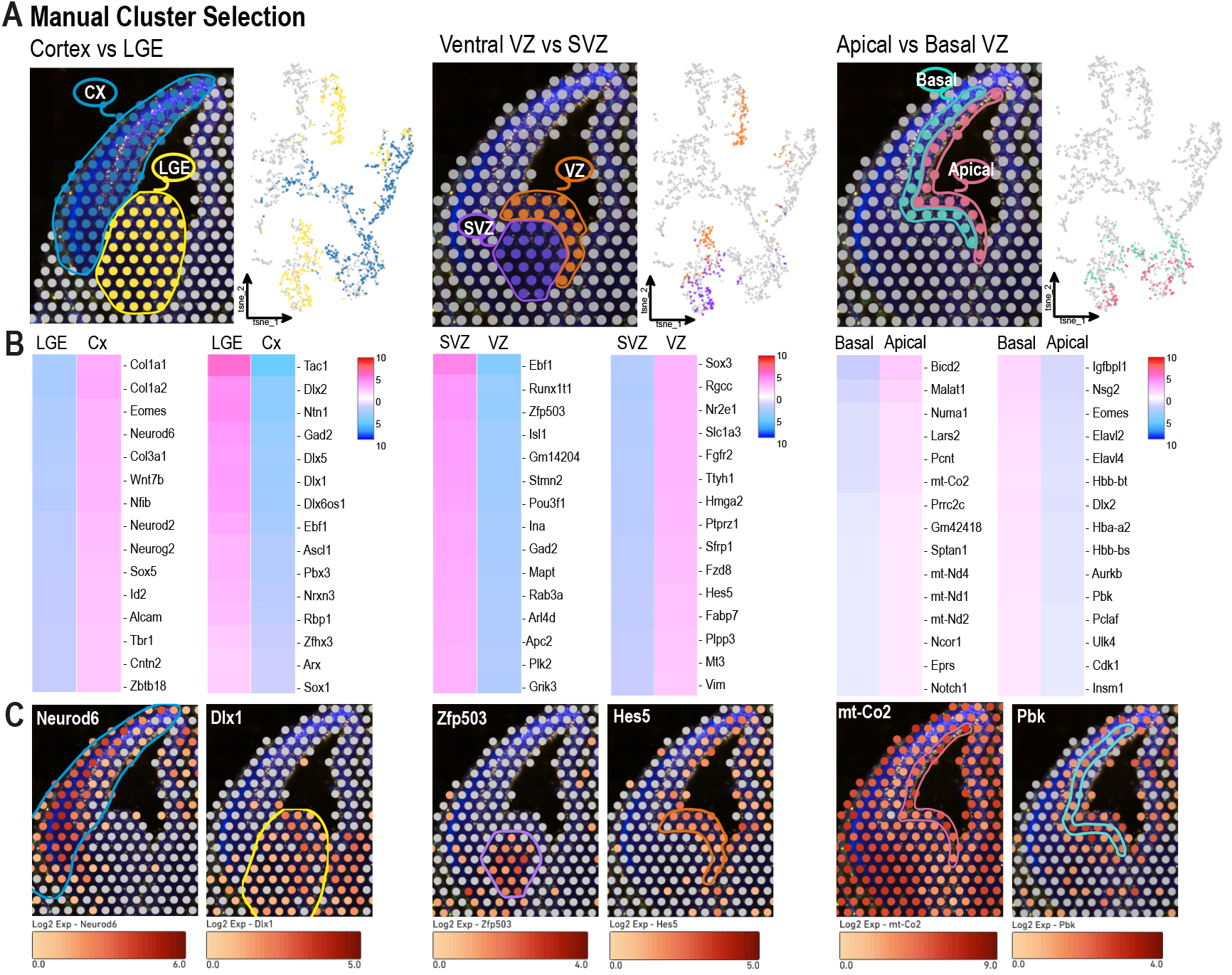
Spatial transcriptomics enables manual cluster selection and comparison. A. Examples of customized cluster selection. The indicated areas were manually selected on the spatial map. On the right, the t-SNE plots for each selected area. Cx: Cortex, LGE: Lateral Ganglionic Eminence, VZ: Ventricular Zone, SVZ: Subventricular Zone. B. Local gene expression comparisons were performed between selected clusters. The panel shows gene expression heatmaps of the top 15 differentially-expressed genes per manual cluster defined in A. C. Spatial expression of representative genes per manual cluster defined in A.

Subsequently, we restricted the comparison to a smaller brain area in the LGE, manually selecting the ventral VZ and SVZ. We defined the limit between the two zones guided by the mitotic layer (PH3+ cells) that marks the beginning of the SVZ (Fig. 3A). These two proliferative areas generate GABAergic neurons. However, while the VZ is mainly populated by NSCs, the SVZ contains intermediate progenitors and postmitotic neurons (Turrero García and Harwell, 2017) (see scheme in Fig. 2A). As the gene expression comparison focused exclusively on the selected clusters excluding data from other spots, we were able to identify subtle differences within the LGE. In addition to expected DEGs, we picked up other genes enriched in the LGE SVZ and VZ, for instance, *Runx1t1* and *Nek6*, respectively (Fig. 3B and Table S2).

Finally, the immunohistochemical images allowed us to further narrowed down the area of interest and we conducted a subregion analysis comparing apical and basal VZ from both dorsal and ventral areas (Fig. 3A). We defined the apical VZ as the mitotic cell layer (PH3+ cells) lining the ventricle and the basal VZ as the adjacent area reaching the boundary between the VZ and SVZ. These subregions host mainly radial glia cells but also intermediate progenitors, particularly the basal zone. Of note, radial glia cells undergo interkinetic nuclear migration, whereby the mitotic phase takes place at the most apical side and the S-phase at the basal VZ (Takahashi et al., 1993). Thus, we expected to uncover transcriptional changes inherent to different cellular states within a single cell type. Interestingly, we found a significant representation of mitochondrial respiration genes in the apical layer, suggesting a metabolic signature of mitotic RG cells. In addition to intermediate progenitor genes (e.g., *Eomes*), we found other transcriptional and post-transcriptional regulators, such as *Elav2/4* enriched in the basal area (Fig. 3B, Table S2). Logically, the number of DEGs exhibiting statistically significant differences in these closely associated areas was smaller compared to the other cases. Finally, to confirm the validity of the analysis, we show the spatial expression of several genes enriched in each cluster (Fig. 3C). Additional cluster types can be defined by the reader by downloading the complete CLoupe file in GEO *(accession number will be provided soon)*.

### Integration of single-cell and spatial transcriptomics data

The advent of scRNAseq technology has revolutionized our ability to study gene expression profiles and understand cell heterogeneity. Nowadays, scRNAseq is a widespread methodology performed in most molecular biology labs and has already been applied to a plethora of biological systems. Thus, the amount of single-cell expression generated data has grown exponentially over the last years and will likely continue expanding in the future (Svensson et al., 2020). Although scRNAseq offers single-cell transcriptomics resolution at a large scale, it lacks spatial information. To provide tissue context to scRNAseq datasets, single-cell data from embryonic brains can be integrated into the ST map. We offer the possibility of mapping embryonic brain scRNAseq data from other labs into our ST maps. As an example, we integrated publicly available scRNAseq data from dorsal and ventral E13.5 embryonic brains (La Manno et al., 2021) with our ST data. The datasets from the two brain areas were merged and their reads were normalized prior subsequent clustering by using the R package Seurat. Next, we mapped the identified clusters onto our ST map by using an anchor-based integration method included in this tool (Fig. 4A). Most clusters corresponded to a defined brain region, offering a straightaway method to assign cluster identity in the scRNAseq dataset (Fig. 4B). For example, cluster 2 mapped to the cortical SVZ and preplate; thus, it is most likely composed of glutamatergic intermediate progenitors and neurons. On the other hand, cluster 4, that mapped into the VZ, is presumably constituted by NSCs. Indeed, molecular signatures of each cluster obtained from the scRNAseq data analysis confirmed the cell type identity (Table S3). The few clusters that did not show a clear spatial distribution are composed of cell types sparsely distributed in the tissue, such as microglia (cluster 9, data not shown) or blood cells (cluster 10). In conclusion, Spatial and single-cell data integration represents an alternative and accurate method for cell type classification.

**Figure 4.**
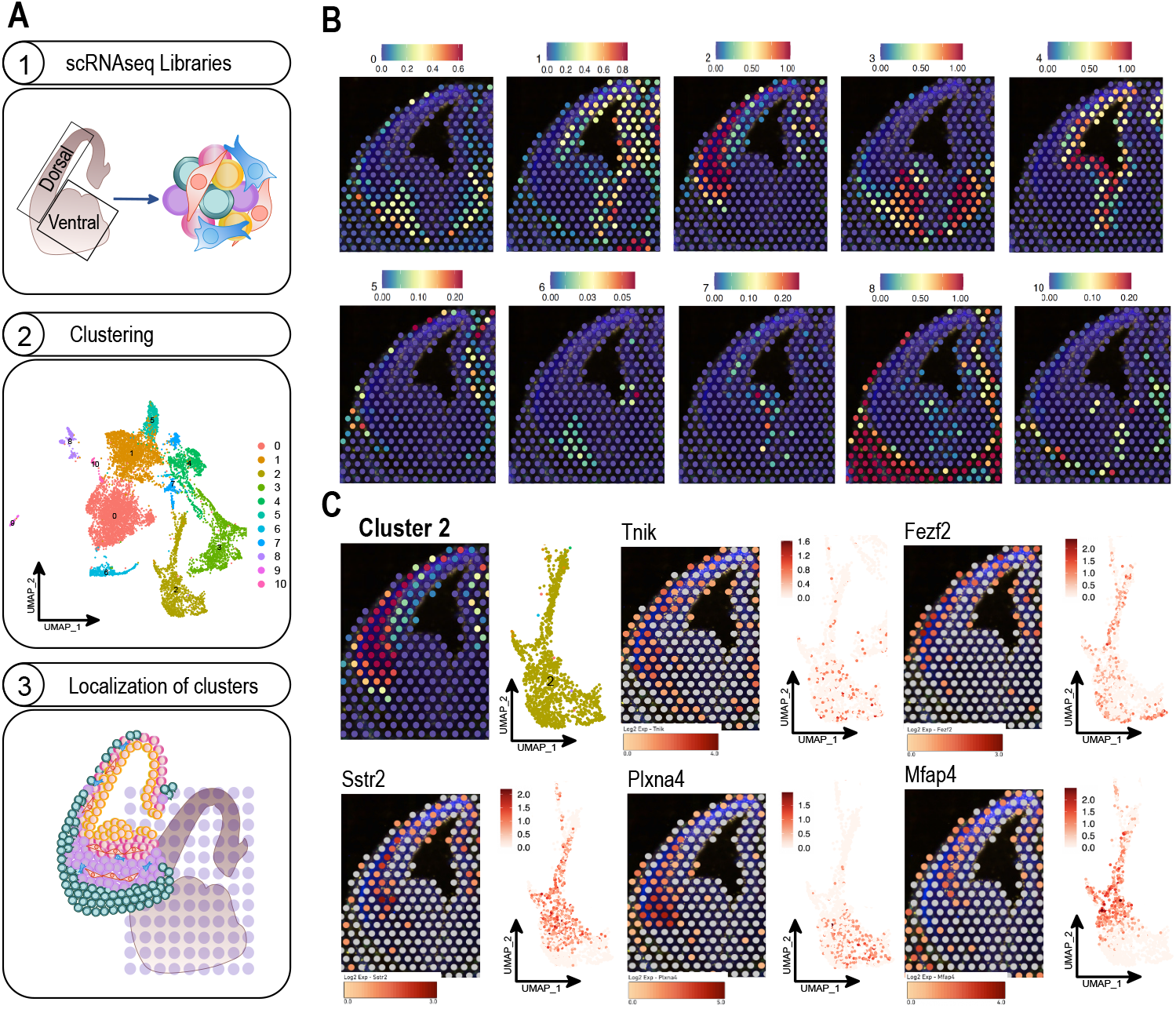
Integration of single-cell and spatial transcriptomics data. **A**. Schematic overview of the workflow combining scRNA-seq and ST data. (1) scRNA-seq was performed from E13.5 mouse brains. (2) Unsupervised data clustering identified 11 groups (3). The annotations from the scRNAseq analysis were transferred to the Spatial Transcriptomics data query by following probabilistic integration methods. **B**. scRNAseq data of E13.5 dorsal and ventral brain were obtained from La Manno et al., 2021. Clusters shown in UMAP plot in A were mapped onto our Spatial Transcriptomic map using the R package Seurat. The colour code represents the prediction scores obtained per spot and cluster. Only the annotations from cluster 9 could not be integrated into any spot in the query ST set. **C**. Spatial visualization of top upregulated genes within cluster 2 (in B) showing a clear subregional expression in the dorsal telencephalon.

scRNAseq data allows to sort cell types according to their molecular signatures via clustering analysis. However, many scRNAseq group-defining genes are not evenly expressed within the cluster. By interrogating the ST map, we elucidated whether high-expressing cells for certain DEGs were sparsely distributed within the tissue or confined to specific subregions (Fig. 4C). Using cluster 2 as an example, we found top upregulated genes with clear subregional expression within the dorsal telencephalon, probably reflecting different neuronal maturational states. Furthermore, by visualizing protein expression patterns for PH3 and CD31, our ST map provides information about the proximity of high-expressing cells for the gene of interest to actively mitotic areas or blood vessels.

Recently, scRNAseq dataset processing by specific bioinformatics tools, such as CellChat and CellPhone, has been useful when searching for cell-cell molecular interactions (Efremova et al., 2020; Jin et al., 2021; Vento-Tormo et al., 2018). However, this analysis does not evaluate physical proximity, a crucial requirement for cell communication. Combined scRNAseq and ST provide in-depth transcriptomic information of single cells in their tissue context and can help elucidate cell-cell molecular partners. In conclusion, ST approach considerably improves the characterization of brain tissue architecture, and details at the single-cell level can be accomplished by combining ST with scRNAseq datasets. Latest advances in molecular biology and bioinformatics open up a new way to study the impact of tissue organization and cell-cell interaction during neurogenesis.

## MATERIALS AND METHODS

### Mouse

C57BL/6N (Charles River) were bred in the DKFZ animal facility (Zentrales Tierlabor, ZTL). The day of the vaginal plug was considered embryonic day (E) 0.5. Embryos were collected at E13.5. Mice were housed in standard housing conditions following the German Animal Welfare Act regulations. All animal experiments were approved by the local governing guidelines (Regierungspräsidium Karlsruhe, Germany).

### Tissue preparation, immunofluorescence staining and imaging for spatial transcriptomics

Pregnant mice were anesthetized via isoflurane inhalation and sacrificed by cervical dislocation. E13.5 mouse heads were snap-frozen on dry ice immediately after dissection and stored at −80°C until use. Subsequently, the fresh frozen brains were embedded in ice-cold Tissue-Tek O.C.T. (Leica) and sectioned by a cryo-microtome (Leica CM1950). Ten μm thick cryosections were collected on Visium Spatial Gene Expression slides (Visium Spatial Gene Expression Slide & Reagents Kit, 10xGenomics, 1000187). Each section was collected into a 6,5 × 6,5 mm capture area surrounded by a fiducial frame of 8 × 8 mm. A total of 4 sections from 4 different embryonic brains were used. After collection, brain slices were fixed in −20°C methanol and stained following the manufacturer’s instructions (10x Genomics, https://www.10xgenomics.com/products/spatial-gene-expression). The following primary antibodies were used as markers: anti-phosphorylated histone H3 (PH3 mouse, 1:500, Abcam, ab14955), anti-Nestin (chicken, 1:200, NOVUS, NB100-1604), anti-CD31 (rat 1:200, BD Pharmingen, 550274), and DAPI (1:1000, Invitrogen, D1306). Brightfield imaging - to capture the section within the fiducial frame - and immunofluorescent imaging - to detect protein expression - were obtained with Tissue scanner Olympus VS200. Both types of pictures were subsequently used for spatial mapping and tissue reconstruction.

### Visium spatial gene expression and library construction

After imaging, the tissue was then processed for permeabilization and cDNA synthesis as described in 10x Genomics Visium Spatial Gene Expression Kit (https://www.10xgenomics.com/support/spatial-gene-expression-fresh-frozen/documentation/steps/library-construction/visium-spatial-gene-expression-reagent-kits-user-guide). Briefly, the tissue was permeabilized in order to release mRNA from the cells that subsequently was bound with spatially barcoded oligonucleotides attached in the capture area. A tissue optimization step was performed beforehand and a permeabilization time of 10 min was established (https://www.10xgenomics.com/support/spatial-gene-expression-fresh-frozen/documentation/steps/tissue-optimization/visium-spatial-tissue-optimization-reagents-kits-user-guide). Barcoded cDNA was then obtained from captured mRNA and subjected to cDNA amplification and library construction. The Visium Spatial Gene Expression library was sequenced by Illumina NovaSeq 6000.

### Visium spatial data visualization and gene expression analysis

Reconstruction, visualization, and spatial gene expression analysis were performed by 10x Space Ranger Software (version 1.3.0) and 10x Loupe Browser (version 5.0) (https://support.10xgenomics.com/spatial-gene-expression/software/pipelines/latest/what-is-space-ranger). Loupe Browser was used to establish a manual fiducial alignment and therefore an accurate tissue boundary selection on every image. The resulting JSON file containing the coordinates of every spot labelled as part of the tissue and their corresponding scaling factors that converts pixel positions from full-resolution to downsampled images were used in combination with its corresponding image in TIFF format and the FASTQ files obtained after sequencing to map every read to the mouse reference genome (mm10-2020-A) by using *spaceranger count*. The resulting four output CLoupe files were merged into one single dataset by using *spaceranger aggr*. This new file was then imported into Loupe Browser for gene expression analysis and manual clustering. To increase the overall resolution of the spatial transcriptomics atlas obtained in the merged CLoupe file, the filtered feature-barcode matrixes of the four samples obtained as a part of the *spaceranger count* output were processed individually using the R package *Seurat* (version 4.3). After normalizing the reads of each replicate individually using the *SCTransform* function, the four Seurat objects were merged. Potential batch effects were removed using *Harmony* and clustering was then performed (16 dimensions, resolution parameter set in 0.5) to get the same number of clusters in all the samples. The processed Seurat object was then split and the list of barcodes containing their corresponding cluster identities was obtained for every single replicate in CSV format. The resulting concatenated CSV file was sent finally as an input into Loupe Browser to give more accurate identities to the clusters identified initially after executing Space Ranger. The final CLoupe file can be downloaded from GEO *(accession number will be provided soon)*.

### scRNAseq and spatial transcriptomics data integration

In order to perform scRNAseq and ST data integration, scRNAseq data of E13.5 dorsal and ventral brain from La Manno et al. were used (La Manno et al., 2021). The BAM files containing the mapped reads of both scRNAseq datasets were downloaded and their feature-barcode matrixes were retrieved by using *dropEst* (version 0.8.6). Filtering, clustering and integration of scRNAseq data onto ST brain map was carried out by using *Seurat*. Briefly, both scRNAseq datasets were converted into Seurat objects and pre-processed individually. Genes expressed in less than 3 cells were filtered out. Cells expressing less than 200 genes, cells expressing more than 2500 genes and cells showing a mitochondrial expression ratio above 5% of the total gene expression were removed to minimize the number of empty droplets, potential doublets and dying cells in the final dataset, respectively. Next, both datasets were merged into one Seurat object and the raw reads were normalized by using the *SCTransform* function. The difference between S-phase and G2/M-phase scores, calculated by using the function *CellCycleScoring*, was regressed out during data normalization. Then, clustering was made by following the standard workflow (resolution parameter set in 0.2). Integration was performed according to the guidelines specified in Seurat into our already processed spatial transcriptomics data (see previous section).

## Acknowledgements

We thank the DKFZ Light Microscopy core facility, the DKFZ Single-Cell Open Lab (scOpenLab) and the DKFZ GPCF High Throughput Sequencing Facility for assistance with sample preparation and Spatial Transcriptomics experiment. We thank Melanie Lutz for excellent technical assistance.

## Author contributions

Conceptualization: B.D.M., J.A.; Methodology: B.D.M., J.V.; Software: J.V.; Validation: B.D.M., J.V. J.A.; Formal analysis: B.D.M., J.V., J.A.; Investigation: B.D.M., J.A.; Resources: L.C., H.M., J.A.; Data curation: B.D.M., J.V., J.A.; Writing - original draft: B.D.M., J.A; Writing - review & editing: B.D.M., J.V., H.M., L.C., J.A.; Visualization: B.D.M., J.V. L.C., J.A.; Supervision: L.C., J.A.; Funding acquisition: L.C., J.A.

## Data availability

All data can be found within the article, its supplementary information, and the following websites: STomicsDB and GEO *(accession numbers will be provided soon)*.

